# Reassessing the “sexy” X hypothesis reveals distinct cellular and disease signatures of X-linked genes in male sexual differentiation and reproduction

**DOI:** 10.64898/2026.02.03.703532

**Authors:** Katia Ancelin, Praveenan Somasundaram, Rafael Galupa

## Abstract

The human X chromosome (chrX) has long been proposed to be enriched in genes involved in brain development, reproduction and sexual differentiation, earning the epithet of “smart and sexy” chromosome. Many studies have confirmed that the chrX is indeed “smart”, but it is less clear whether “sexy” still holds true. Here, we investigated human X-linked genes with respect to their expression levels across tissues and cell types, their annotated functions, and their association with monogenic disorders affecting sexual differentiation and reproduction (SDR). We confirm that many brain and sex-specific tissues show high X-to-autosome (X:A) expression ratios; however, paradoxically, some of the corresponding main cell types, such as neurons or prostatic or endometrial cells, show X:A ratios below the average. In contrast, cell types such as blood and vascular cells have elevated X:A ratios, suggesting that they contribute to the high ratios observed at the tissue bulk level. Analysing the most highly expressed genes across tissues and cell types, we observed a significant enrichment in chrX genes for testes, spermatogonia and Sertoli cells. Interestingly, monogenic disorders with SDR phenotypes showed a significant enrichment of chrX genes only when related to male-specific conditions. Our study thus supports a male-biased view of the "sexy" human chrX, whereby it is particularly enriched in genes highly expressed in testicular cell types and in genes involved in male reproductive monogenic disorders. Intriguingly, these genes overlap very little, suggesting an important disconnect between expression levels and clinical associations, at least on the X chromosome.

## INTRODUCTION

More than two decades ago, the human X chromosome (chrX), which harbours an estimated total of 839 protein-coding genes, earned the epithet "smart and sexy" due to its atypical gene content involved in neuronal functions, sex determination, and reproduction (1,2). Over the years, different studies have supported the adjective “smart” (3–5), including a recent systematic analysis that revealed an enrichment for the human chrX (over autosomes) in genes implicated in disorders related to cognition, language and epilepsy (6). Less attention has been given to whether the adjective “sexy” is still applicable, despite fertility problems associated with X-linked aneuploidies and the societal impact posed by increasing infertility worldwide (7–9).

Originally, Graves and colleagues classified the human X chromosome as “sexy” (1,2) based on two studies that reported an overrepresentation on the human X of genes related to sexual differentiation and reproduction (SDR). In 1999, before the first draft sequence of the human genome, Saifi and Chandra searched medical databases to identify genes with mutations affecting SDR with no explicit distinction between sexes, and reported that there was a significant enrichment of approximately three-fold for chrX genes (10). In parallel, while analysing 25 genes expressed in mouse spermatogonia but not in somatic tissues, David Page’s lab found that 10 were X-linked, a number that is at least five times higher than if there was a random distribution across all chromosomes (11). Since then, other studies have analysed the human and mouse chrX gene content in relation to SDR; in **Table 1** we have summarised their findings. Most studies point to the chrX being enriched in genes expressed in the male gonads; this has been proposed to be dominated by the cancer/testis antigen (CTA) genes, which are highly enriched on the chrX (12,13). CTA genes are characterised by expression in several cancer types, while their expression in normal cells is solely or predominantly in testis (12). However, the function of these genes in germ cell development and male reproduction is largely unknown, so the functional meaning of their expression is still unclear.

Different approaches have been used across studies to evaluate the chrX gene content in relation to SDR, which can help explain the different conclusions reached: in some studies, authors evaluated SDR content in terms of genes with mutations affecting SDR (10), while others in terms of chrX gene expression in reproductive organs – either regarding levels of expression (5,11,14–16), or their representation among the most highly expressed genes in each tissue or among tissue-specific genes (17–19). Given that these studies were conducted more than a decade ago, we sought to systematically reassess the longstanding hypothesis of the “sexy” X. Based on current resources for transcriptomics, functional annotations and clinical genetics, we re-evaluated human X-linked genes across tissue- and cell-type-specific expression datasets, SDR-related functional annotations and associations with monogenic SDR disorders.

## RESULTS

### High X-to-autosome expression ratios are not restricted to brain-specific or sex-specific tissues and cell types

To analyse gene expression data, we used the Human Protein Atlas (HPA) database, which combines curated human transcriptomic and proteomic data, including from the GTEx portal (20,21), and downloaded bulk “consensus” transcriptomic data from 50 different tissues, and aggregated single-cell transcriptomic data for 154 cell types (see Materials and Methods). Data for non-sex-specific tissues and single cells is from male and female individuals combined. Expression ratios between the chrX and autosomes (X:A ratio) have been used as a proxy for assessing dosage compensation, and have revealed high values for brain tissues (5). To explore this further, we determined X:A ratios across different tissues and cell types (see Materials and Methods section for more details). Given how lowly expressed genes can skew such types of analyses (16,22), and to define a threshold for “expressed” genes, we first plotted the distribution of log-transformed normalised Transcripts Per Million (nTPM) or normalised Counts per Million (nCPM) values for the entire tissue or single-cell datasets, respectively (**Fig. S1A-B, Supplementary data sheet 13**). In both cases, a typical bimodal distribution was observed (23), and we filtered out genes with less than three nTPM/nCPM (**Fig. S1A-B**). This left us with 54% of each global dataset, including ∼18.500 genes (91-92% of the total). We calculated the overall average X:A expression ratio, which was ∼0.93 for the tissue dataset, in line with previous studies (e.g., 0.94 in (5)), and ∼0.87 for the single-cell dataset. This supports the notion of nearly dosage compensation between X-linked and autosomal expression, given that there is only one active allele for X-linked genes in either males or females but two active autosomal alleles (22).

For nine adult brain tissue datasets, we observed the expected X:A expression ratio higher than average (∼1.04, purple dotted line, **Fig. S1C)**. However, when analysing single-cell datasets from brain-related cell types, we observed an X:A expression ratio of ∼0.73, below the average; this was the case, for instance, for the pituitary-gland cell types (**Fig. 1A**) or most brain-specific cell types (**Fig. 1B**). We reasoned that this surprising discrepancy between tissue data and single-cell data could perhaps reflect the contribution of other cell types: brain tissues also contain fibroblasts, immune cells, mural and endothelial cells, among others. Interestingly, all these cell types show X:A expression ratios above the average (**Fig. S1D**), suggesting that they could indeed contribute to skew the brain X:A ratios to higher values. Thus, the reported “high X:A expression ratios” in the brain could in fact be due to cell types that are not brain-specific. We do note that the aggregated single-cell data from HPA is not always tissue-specific; for instance, the dataset corresponding to “fibroblasts” includes fibroblasts from different tissues and not just from the brain.

**Figure 1.**
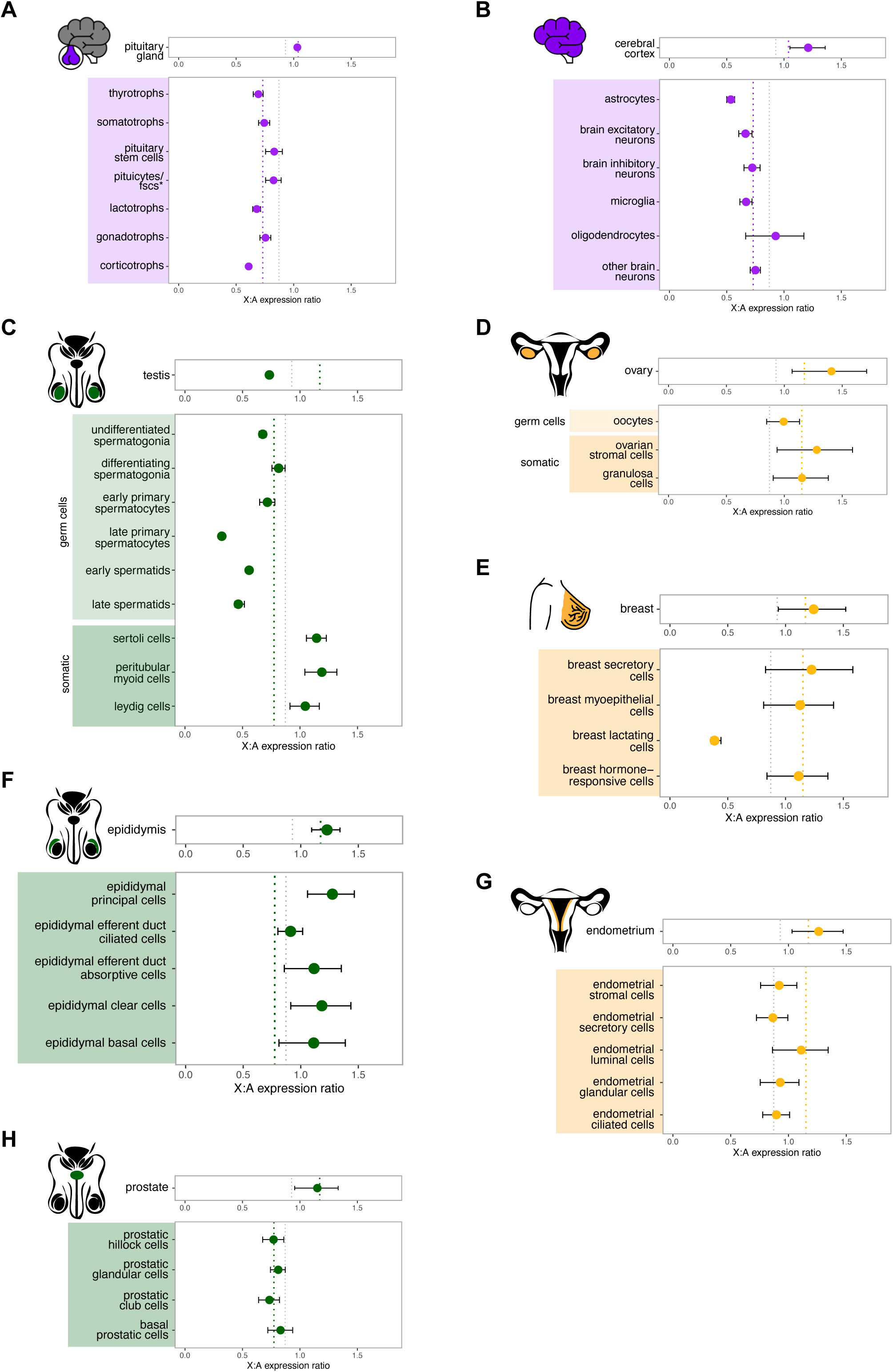
X-to-autosome expression ratios in brain-related and sex-specific tissues and cell types. Each panel shows, at the top, the X:autosome expression ratio for a given tissue: pituitary gland (A), cerebral cortex (B), testis (C), ovary (D), breast (E), epididymis (F), endometrium (G) and prostate (H); at the bottom, each panel shows X:autosome expression ratios for cell types specific to each tissue. Each point represents the mean X:autosome expression ratio (± s.e.m.) for a given tissue or cell type listed on the left. Coloured dotted lines represent the average X:autosome expression ratio for each of the groups of tissues or cell types analysed. Grey dotted lines represent the overall average X:autosome expression ratio for either the tissues dataset (∼0.93) or the single-cells datasets (∼0.87).

In sex-specific tissues (11 tissues) we observed an even higher mean X:A expression ratio (∼1.17, red dotted line, **Fig. S1C**). Of note, the testes were an exception, showing an overall low average X:A expression ratio (∼0.75); such a trend has been reported previously (5). This is likely related to the lower expression of X-linked genes in late spermatogenesis (18) due to the meiosis-associated silencing of the X chromosome in male germ cells (14). Indeed, analysis of average X:A expression ratios in testicular cell types revealed lower values for late-stage germ cells (**Fig. 1C**; e.g., late spermatocytes, spermatids). In contrast, testicular somatic cells show high X:A ratios (**Fig. 1C**). The relative composition of the testis by these different cell types, with germ cells largely outnumbering somatic cells (24,25), probably explains the low X:A ratio observed at the tissue level.

The ovary shows a high X:A expression ratio, which is shared by the ovarian somatic cells (granulosa and stromal cells; **Fig. 1D**). Oocytes (the ovarian germ cells) show lower X:A expression ratio, in line with previous microarray data (5). Other sex-specific tissues, like the epididymis and the mammary gland, also show good concordance between tissue and cell-type X:A ratios (**Fig. 1E-F**), while the prostate and the endometrium show higher tissue X:A expression ratios than their corresponding cell types (**Fig. 1G-H**). Given that both prostate and endometrium are highly vascularised organs, the higher X:A ratios at the tissue level could be due to contribution from blood and endothelial cells (**Fig. S1D**), as hypothesised above for the brain.

While we confirm high X:A ratios for brain-related and most sex-specific tissues, our analyses also reveal that in many of these tissues the high values are not associated with brain-specific or sex-specific cell types, and suggest they rather associate with “pervasive” cell types such as blood and vascular cells.

### X-linked genes are enriched among the most highly expressed genes in human testis and some of its cell types

Previous studies have analysed the proportion of X-linked genes among highly expressed genes in sex-specific tissues, with mixed results (**Table 1**). To revisit this, we downloaded the ‘elevated expression’ transcriptomic data curated from the HPA and GTEx databases (20,21), which includes the most highly expressed genes across 36 different tissues and 80 different cell types (26); we again classified them as ‘brain-related’, ‘male-specific’, ‘female-specific’ and ‘others’ (see Materials and Methods section for more details). To determine whether chrX genes were enriched or depleted among the ‘elevated expression’ datasets, we compared the *observed* number of protein-coding genes per tissue, for each chromosome, with the *expected* number, calculated based on the total number of protein-coding genes for each chromosome. We present the results for chrX along with three autosomes – chr5, chr7 and chr16 – chosen for their similar number of protein-coding genes (chrX: 839; chr5: 813; chr7: 888; chr16: 815). While chrX genes were almost always observed at lower frequency than expected in ‘other tissues’ (**Fig. S2A**), in several sex-specific tissues they were observed at higher frequency than expected (testis: 2.17x; ovary: 2.83x; epididymis: 2.15x; endometrium: 2.31x; **Fig. 2A**, middle and bottom). This was only statistically significant for the testis (**Fig. 2A**, middle), perhaps due to the considerably higher number of “elevated expression” genes for this tissue compared to the others (testis: 1992; ovary: 178; epididymis: 96; endometrium: 89). It is interesting to note an enrichment of chrX genes in testis – among its most highly expressed genes – despite a lower X:autosome expression ratio, which considers levels of expression for all expressed genes. Interestingly as well, for brain tissue there is no significant enrichment of chrX genes among its highly expressed genes (1.24x; **Fig. 2A**, top), despite a high X:autosome expression ratio.

**Figure 2.**
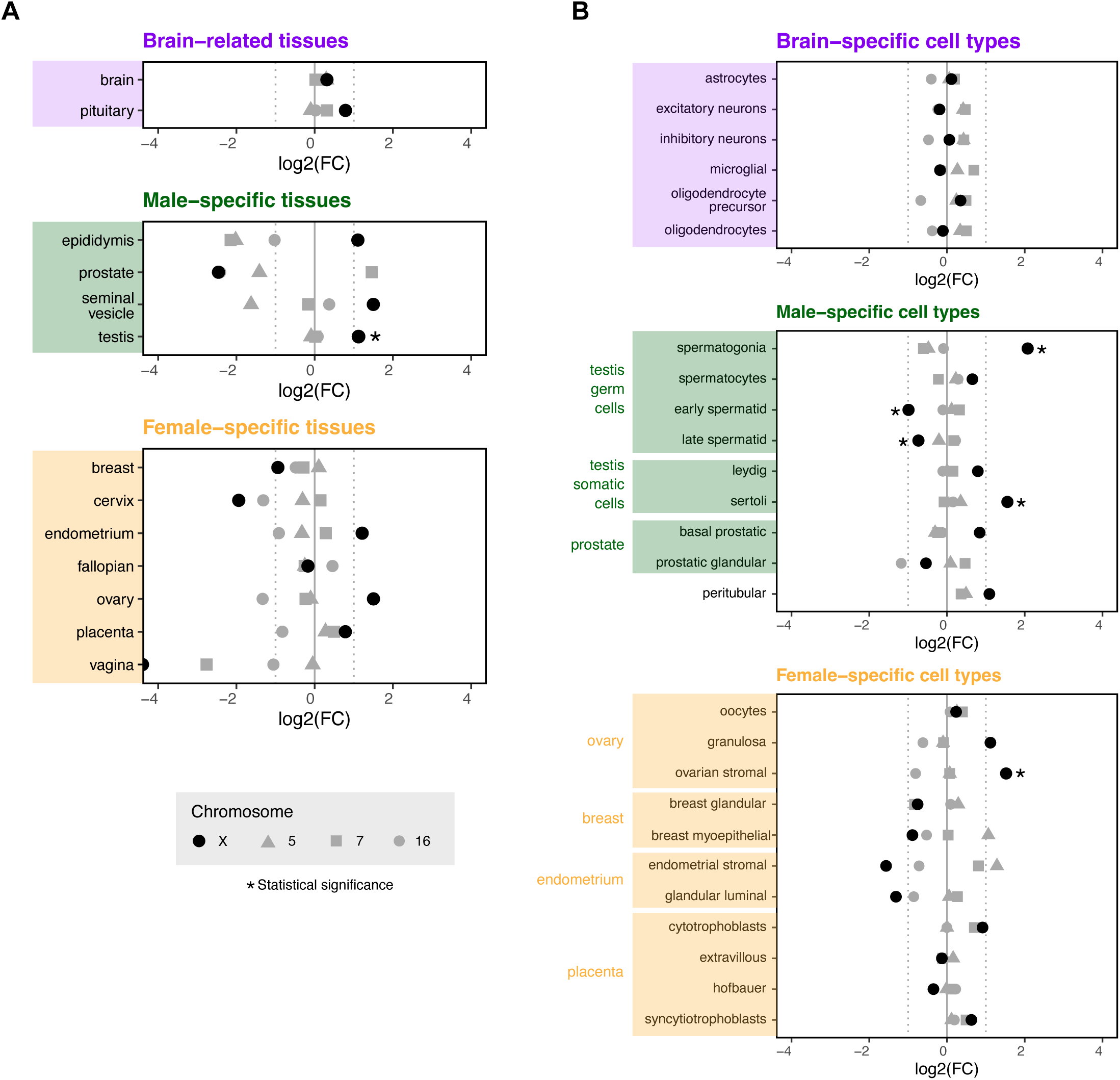
X-linked genes are enriched among highly expressed genes in testis and in specific testicular cell types. Dot plots for log2-transformed ratio (fold-change, FC) between observed and expected frequency of chrX genes among the “elevated” group of genes for tissues (A) or cell types (B) as downloaded from the Human Protein Atlas webpage (version 23.0). For panel A and B, brain-related tissues and cell types are shown at the top; male-specific tissues and cell types are shown in the middle; female-specific tissues and cell types are shown at the bottom. Statistical test: Chi-squared or Fisher’s test (see Materials and Methods); significance (*p*-value<0.05) is represented as a black asterisk next to the respective dot (see Supplementary Data sheet 3 for *p*-values). Multiple testing correction was performed using false discovery rate (FDR).

In concordance with the analysis for the tissues, chrX genes were almost always observed at similar or lower frequency than expected in ‘other cell types’ (**Fig. S2B**). For brain-related cell types, the observed number of chrX genes was close to expected (**Fig. 2B**, top), again in agreement with the tissue analysis. For female-specific cell types, the picture was more heterogeneous (**Fig. 2B**, bottom). Interestingly, for stromal cells, which represent the most prevalent cell population in the ovary, the enrichment of chrX genes was statistically significant, which likely contributes to the enrichment we observed in the ovary (**Fig. 2A**, bottom); a large proportion of the most highly expressed X-linked genes in stromal cells (n=30) and the ovary (n=22) is common (n=13; **Supplementary data sheet 5**). Finally, for male-specific cell types, we observed a high observed-to-expected ratio for somatic cells (Sertoli, Leydig, peritubular and basal prostatic cells; **Fig. 2B**, middle), which is statistically significant for Sertoli cells. For the male germ cells, the trend seems to reflect the dynamics of chrX expression during spermatogenesis (27): in spermatogonia, the undifferentiated germ cells that act as “stem cells”, there is a significant enrichment for chrX genes (**Fig. 2B**, middle); no enrichment is observed for spermatocytes, the “intermediate” cell type; and a significant depletion for chrX genes is observed for early and late spermatids (**Fig. 2B**, middle), the final stages before spermatozoa. As previously stated, this shift from enrichment to depletion is consistent with the known dynamics of sex-chromosome regulation during spermatogenesis, including meiotic sex-chromosome inactivation and subsequent post-meiotic transcriptional constraints (18). Enrichment on the X of genes predominantly expressed in spermatogonia and Sertoli cells was also observed in a recent study that analysed single-nucleus transcriptome data for testes from ten mammalian species (27). Cross-analysis of the sets of chrX genes with “elevated expression” in testis (n=189), spermatogonia (n=97) and Sertoli cells (n=38) show that 40% of “testis genes” overlap with spermatogonia ones, and 10% with genes expressed in Sertoli cells (**Fig. S2C; Supplementary data sheet 5**), suggesting a bigger contribution of the germ cells to the bulk tissue results, consistent with their relative abundance.

We wondered whether CTA genes could be related to the significant enrichment of chrX genes in the testis and its cell types. As stated previously, CTA genes are highly enriched on the chrX and specifically expressed in testis and at high levels (28,29). Indeed, CTA genes represent 39% of chrX genes with “elevated expression” in the testis dataset (**Supplementary data sheet 5**), which is ∼four-fold higher than expected by chance, since CTA genes represent ∼10% of the X-linked genes (34). For spermatogonia and Sertoli cells, CTA genes represent, respectively, 43% and 8% of the chrX genes with “elevated expression”, consistent with the overlap observed between spermatogonia and testis and the contribution of the germ cells to the whole-tissue results. When CTA genes were removed from the analysis, the enrichment of chrX genes among the most highly expressed genes in human testis was lost (**Supplementary data sheet 2**); sparser data precluded the same type of analysis for the single-cell datasets.

In summary, these analyses indicate that enrichment of X-linked genes among the most highly expressed genes is not a general feature across tissues or cell types, but is instead highly context-dependent. In the testis, the enrichment observed seems to be largely driven by a subset of testis-specific CTA genes, with contribution from both somatic and early germ-cell populations, while later stages of spermatogenesis show depletion of X-linked genes. Given that the functions of CTA genes are largely unknown, it remains unclear what is the significance of their high expression for the male gonads and male reproductive biology.

### No particular association between X-linked genes and SDR-related annotated functions

We next turned to g:Profiler to perform functional enrichment analysis (also known as over-representation analysis or gene set enrichment analysis) and probe whether chrX genes are significantly associated with SDR-related functions. chrX genes, whether CTA or not, are significantly associated with several molecular functions and biological processes, but none directly related to SDR (**Fig. 3A; Supplementary data sheet 6-8**). We performed the inverse analysis and used AmiGO 2 (31–33), a web-based set of tools for searching and browsing the Gene Ontology database, to retrieve all genes with annotated SDR-related functions (n=1761), including “gonadal development” and “sex differentiation” (full list in **Fig. 3C**). For comparison, we also retrieved all genes associated with “brain development” (n=728) and “heart development” (n=569). In line with the established role of the chrY in sex determination and differentiation, chrY genes were significantly enriched among genes related to “sex differentiation”, “gonad development” and “male gamete generation” (**Fig. 3C**). ChrX genes, however, were not enriched among any of the SDR-related classes of functional annotations. Similarly, genes annotated with “brain development” were not enriched for chrX genes, despite previous observations of an enrichment of chrX genes associated with brain-related disorders (6). This discrepancy might reflect distinct biological information captured by functional annotations compared with disorder-based annotations (34,35). These results illustrate that the extent to which the X chromosome can be considered functionally specialized depends strongly on the type of evidence considered.

**Figure 3.**
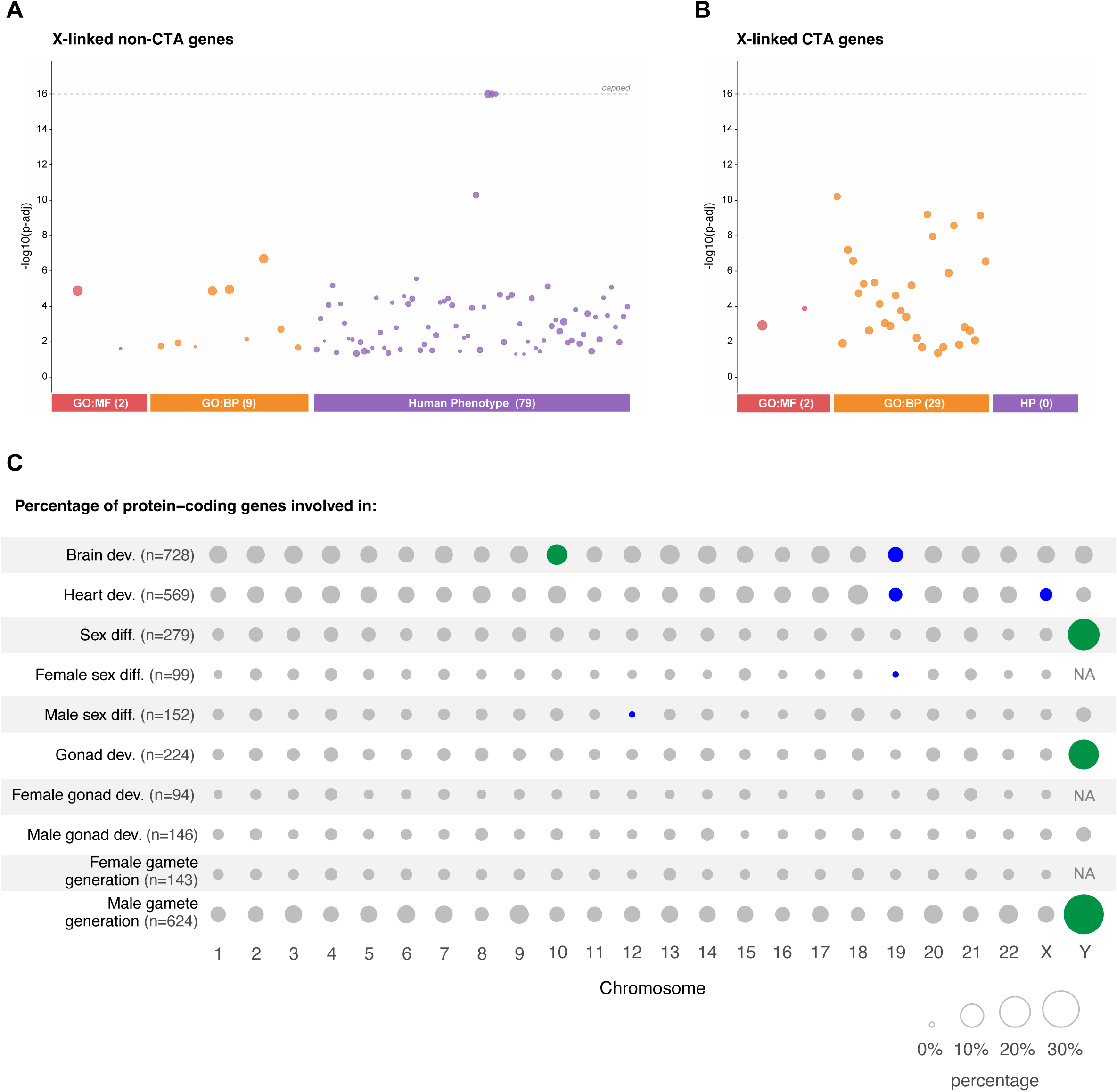
No particular association between X-linked genes and SDR-related annotated functions. (A-B) Manhattan plots for non-CTA X-linked genes (A) and X-linked CTA genes (B) display significantly enriched terms from multiple annotation databases (via g:Profiler), including Gene Ontology Biological Process (GO:BP), Molecular Function (GO:MF) and Human Phenotype (HP). Each circle represents one enriched term, with the y-axis indicating enrichment significance as −log10(adjusted p-value). Circle size is proportional to the number of query genes associated with the enriched term. (C) Bubble plot showing the proportion of genes for each chromosome among genes with annotated functions listed on the left. Green colour represents statistically significant enrichment, while blue colour represents statistically significant depletion (Chi-squared test, *p* value < 0.05). Circle size indicates the percentage of genes involved.

### X-linked genes are enriched among genes associated with monogenic disorders related to male-specific SDR

g:Profiler results also revealed that non-CTA chrX genes are significantly associated with several human disease phenotypes (**Fig. 3A**). As expected, many of these are related to cognition and behaviour, but also craniofacial morphology and genitourinary morphology (**Fig. S3** and **Supplementary data sheet 6-8**). The latter was particularly rich in terms related to male genitalia (e.g., testicular atrophy, hypospadias, micropenis). Based on these results and inspired by the approach conducted by the Depienne’s lab and collaborators (6), which found an enrichment of chrX genes in monogenic disorders related to brain functions, we investigated whether chrX genes could similarly be enriched in SDR-related monogenic disorders. Using the Online Mendelian Inheritance in Man (OMIM) database (https://omim.org/) and its clinical synopses, we built an extensive list of terms (comprised in between 30 and 50 per category) to retrieved genes associated with (i) SDR-related conditions (including infertility) affecting both sexes (n=261; 16 on the chrX, 6.1%); (ii) male-specific SDR-related conditions (n=487; 58 on the chrX, 11.9%); (iii) female-specific SDR-related conditions (n=119; 11 on the chrX, 9.2%); (iv) endocrine-related conditions, affecting one or both sexes (n=133; 10 on the chrX, 7.5%) (**Fig. 4A**, see Materials and Methods and **Supplementary data sheet 9** for full list of terms). Genes on chrX were significantly more frequently associated with male-specific SDR-related disorders (28%) than genes on autosomes (ranging from 8 to 22%) or chrY (17%) (corrected p-value = 2.6 x 10-9; OR = 3.8; **Fig. 4B-C, Supplementary data sheet 10**). No significant enrichment was detected for the other categories (**Fig. 4D-I**).

**Figure 4.**
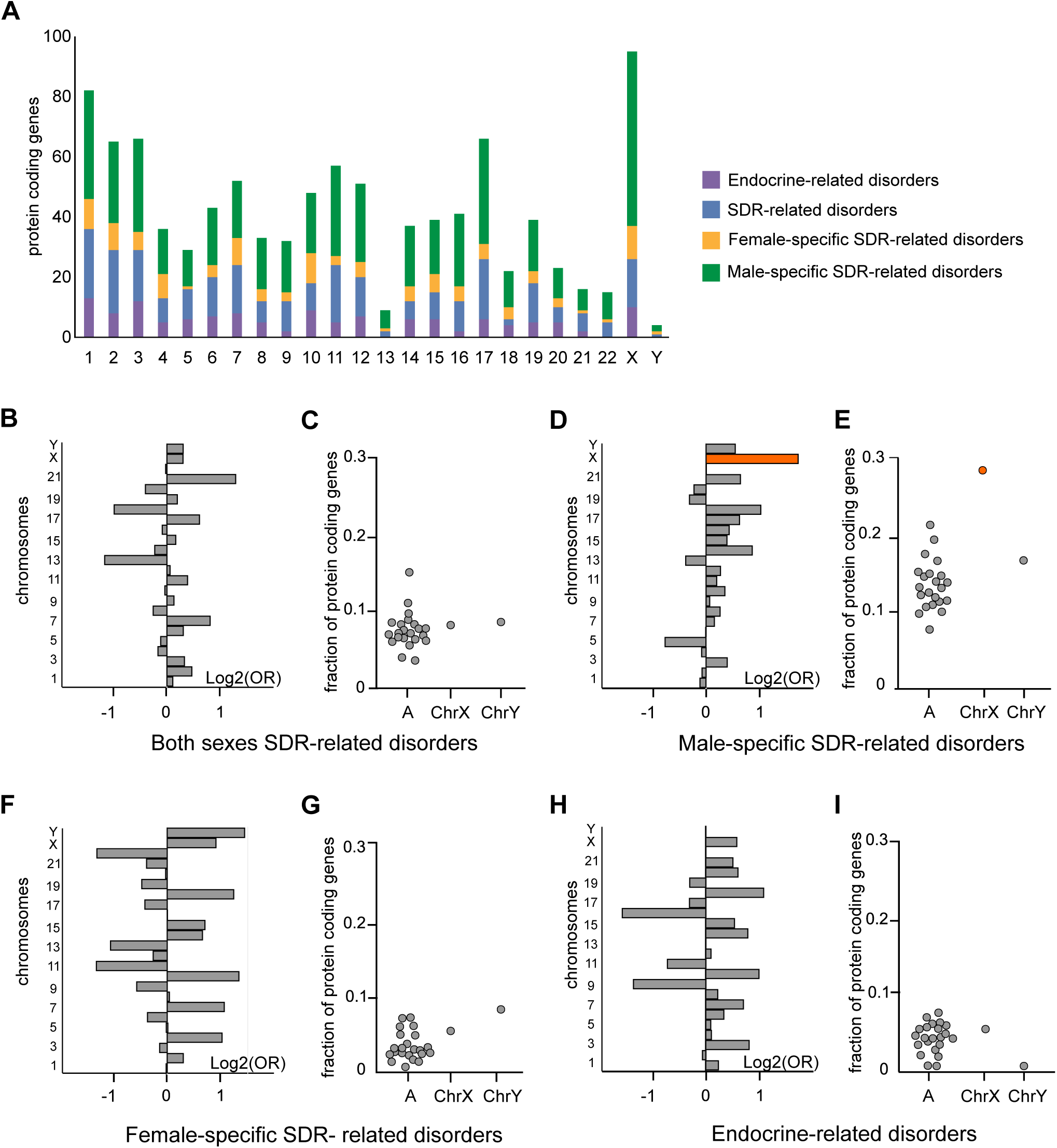
Gene association with SDR-related monogenic disorders. (A) Bar plot showing the number of protein-coding genes per chromosome associated with clinical synopsis data. Categories are indicated in legend; the full list of terms is provided in Supplementary Data sheet 9. (B, D, F and H) Bar plots showing log2-transformed odds ratios for enrichment or depletion of protein-coding genes with SDR-related features. (C, E, G, I) Dot plots showing the fractions of genes per chromosome associated with SDR-related features. Light grey indicates non-significant results, whereas orange indicates statistically significant enrichment (*p* value < 0.01; two-sided Fisher’s exact test followed by Bonferroni correction for multiple testing across chromosomes and phenotypes). Source results for these plots are provided in Supplementary Data sheet 10.

We wondered whether the specific association of chrX genes with male-specific SDR-related disorders could be due to an ascertainment bias related to the fact that men are more often affected by X-linked disorders, given they have a single X chromosome. In line with this, 72% (42/58) of genes associated with male-specific SDR-related disorders are associated with recessive inheritance (**Supplementary data sheet 11**), and 19% (11/58) with dominant inheritance; these trends are inverted for genes associated with female-specific SDR-related disorders: 28% (3/11) of the genes show recessive inheritance, while 64% (7/11) are associated with dominant inheritance. Zechner et al (4) addressed the ascertainment bias for the X chromosome in relation to “smart” genes by comparing the proportion of OMIM entries for chrX and autosomes involved in intellectual disability syndromes (using the term “mental retardation”, meanwhile outdated), with the proportions affecting other malformations, such as skeletal dysplasia. They found that the ascertainment bias is indeed present for all diseases: they observed a ∼2-fold higher frequency of entries for the chrX than for autosomes for diseases such as skeletal dysplasia; however, this frequency was even higher (∼8-fold) for “mental retardation” (4), highlighting an enrichment for chrX genes independent of ascertainment biases. We repeated their analysis and included our SDR-related categories (**Fig. 5A**). Likely due to the more complete picture of the present-day database, which includes almost four times more entries than at the time of the Zechner et al study (4), the ascertainment bias seems overall less pronounced than previously observed (∼0.9-1.5-fold for diseases such as skeletal dysplasia). Yet, we still found a higher enrichment for “mental retardation” (∼2.6-fold). Male-specific SDR-related disorders showed a similar enrichment to “mental retardation” (∼2.5-fold). It thus seems that the male-specific enrichment is present independently of ascertainment biases.

**Figure 5.**
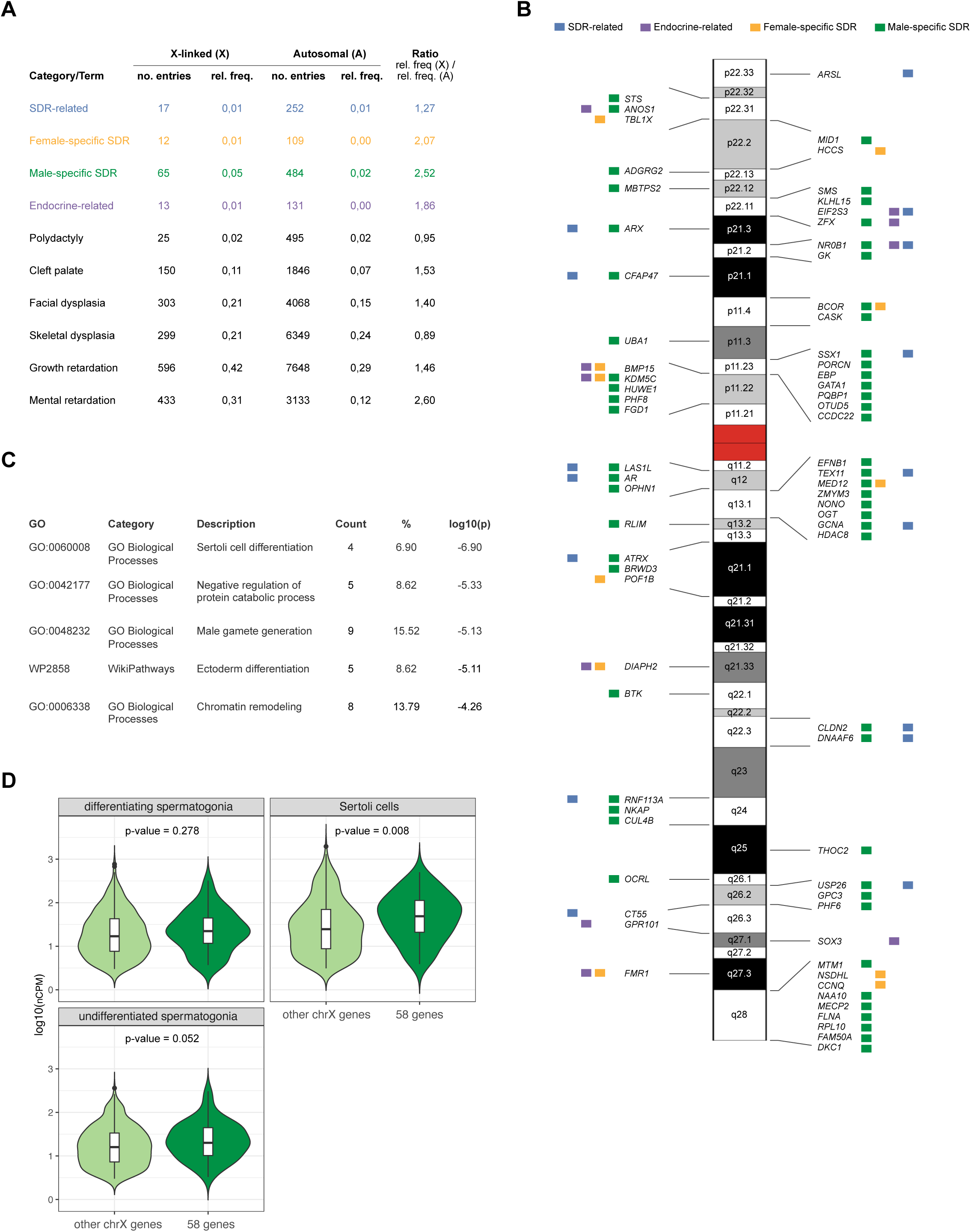
X-linked genes associated with SDR-related monogenic disorders. (A) Number and relative frequency of X chromosomal and autosomal OMIM entries obtained when queried with indicated category/term of disease phenotypes. The frequency was calculated relative to the total number of X chromosomal (n=1410) and autosomal (n=26498) OMIM entries. We kept the outdated term “Mental retardation” for intellectual disability (ID) for direct comparison with the analysis done by Zechner et al. 2001. (B) Localisation on the X chromosome. Squares next to the genes represent association with the studied categories of monogenic disorders. A horizontal line separates genes present in different chromosome bands. Data underlying this scheme can be found in Supplementary Data. The banded chrX was copied from (6). (C) Gene Ontology analysis of genes in the male-specific SDR-related disorders. GO denotes the Gene Ontology identifier for each enriched term. Category specifies the functional class of the pathway or biological process. Description provides the name of biological function or pathway. Count indicates the number of genes from the input list mapping to each term, and % represents the proportion of the total gene list. The log10-transformed enrichment *p*-value, reflects the statistical significance of each term. (D) Violin plots display the distribution of log₁₀(nCPM) expression values for the 58 genes specifically associated with male-specific SDR conditions (dark green) versus all other X-linked genes (light green) in differentiating spermatogonia (top left), Sertoli cells (top right), and undifferentiated spermatogonia (bottom left). Each plot includes a boxplot overlay showing the median and interquartile range. P-values from Wilcoxon rank-sum tests are indicated above each comparison.

In total, 33% of known disease genes on chrX are associated with SDR-related disorders, and they are distributed along the whole chrX (**Fig. 5B; Supplementary data sheet 12**). Genes specifically associated with male-specific SDR conditions represent 28% of known disease genes on chrX (**Fig. 5B**). Since these genes represent the highest proportion of chrX associated with SDR-related disorders, we decided to investigate them further. We ran a gene ontology analysis for these 58 genes (**Fig. 5C**); among the top five most significant terms, we found “Sertoli cell differentiation” (*p* value < 1e-6) and “Male gamete generation” (*p* value < 1e-5) (**Fig. 2F**). Interestingly, a significant proportion of those genes are involved in more general processes such as “Negative regulation of protein catabolic process” (*p* value < 1e-5) or “Chromatin remodeling” (*p* value < 1e-4). Analysis of the spermatogonia and Sertoli cells transcriptomic datasets (HPA database) revealed that in Sertoli cells the 58 genes are significantly more highly expressed than other chrX genes (**Fig. 5D**). Together, these findings suggest that the impact of X-linked genes on male SDR phenotypes might be mediated through cell-type–specific and broad regulatory functions in testicular homeostasis. Of note, only four genes (*DNAAF6*, *GCNA*, *NR0B1*, *SSX1*) are common between the 58 chrX genes associated with male monogenic disorders and the 189 chrX genes among the “elevated expression” dataset in testis, and only one of the 58 genes (*NR0B1)* is among the “elevated expression” genes in Sertoli cells. This further highlights how levels of expression and functional significance are not necessarily correlated; the X-linked gene set among the most highly expressed genes in male-specific cell types and the one underlying male-specific SDR-related disorders is not the same.

## DISCUSSION

We set out to understand whether the human X chromosome, known to be enriched on genes associated with brain functions and diseases, could also be enriched in genes related to reproduction and sexual differentiation. This idea of a “sexy” X chromosome, proposed in 2002 (1), was based on two studies: (i) Wang et al., 2001 found a significant enrichment of X-linked genes among two dozens of genes expressed in mouse spermatogonia (but not in somatic tissues), and (ii) Saifi and Chandra, 1999 found a significant enrichment of X-linked genes among genes with mutations affecting human sexual differentiation and reproduction (10,11). Do we find equivalent results using the latest transcriptomic and clinical datasets? Regarding gene expression in the male gonads, while testis shows the lowest X-to-autosome expression ratio among sex-specific tissues, we found an enrichment of X-linked genes among the most highly expressed genes in human testis, and specifically in spermatogonia and Sertoli cells. This effect, at the tissue level, was ‘dominated’ by CTA genes; among the X-linked genes identified by Wang et al., a third are CTA genes, in line with what we found. Our findings therefore support the original notion of a “sexy” X chromosome. Regarding genes affecting human SDR, our OMIM-based analysis did not reveal an enrichment for chrX genes when considering SDR terms applicable to both sexes. However, we did observe a significant enrichment when considering male-specific SDR terms. Saifi and Chandra (10) used a single list of terms for their analysis, regardless of whether the terms related to conditions that affected only males, only females, or both; the inclusion of the male-only terms might explain why they found a significant enrichment for chrX genes. At this level, our findings partially support the ’sexy’ X hypothesis, pointing to a male-specific bias.

Other studies after the “sexy” X proposal have looked into the relationship between X-linked genes and reproductive tissues (**Table 1**). Lercher et al. (2003) found an enrichment for X-linked genes among prostate-only genes, but not among ovary-only or mammary-only genes; this seems at odds with our findings, but the difference might stem from that fact that the “elevated expression” dataset we used, despite enriched for tissue-specific genes, does not contain only tissue-exclusive genes. In a separate study, Khil et al. (2004) analysed highly expressed genes across mouse tissues using microarray data, and found that chrX genes were enriched in ovaries, but underrepresented in testes. It will be interesting to investigate whether the differences with our conclusions might be species-specific.

We also complemented our analyses with an investigation based on functional annotations. Interestingly, we found no enrichment of chrX genes among genes annotated with SDR-related functions (including male-specific SDR), nor with brain-related functions. The fact that we and others observe an enrichment of chrX genes among genes involved in SDR-related and brain-related monogenic disorders suggests that there is somehow a gap between function and disease, or that genes involved in SDR have pleiotropic roles not captured in the current functional annotations. The origin of this will have to be further investigated; for now, the evidence from monogenic disorders seems of significant enough importance for the X to merit the "smart and sexy" epithet. It is also in line with phenotypes associated with X-chromosome aneuploidies often involving gonadal dysfunctions and infertility symptoms (36,37).

The enrichment of CTA genes on the X chromosome and their specific expression in the male gonads seem to account for the significant enrichment we observed of chrX genes among the most highly expressed genes in the testis. Yet, CTA genes do not account for the specific enrichment of chrX genes among male-specific SDR-related disorders; most genes underlying the male-specific SDR monogenic disorders are not CTA genes (56 out of 58; CTA-family genes are *ARX* and *SSX1*; **Fig. 5**). This further highlights the above-mentioned gap between expression, function and role in disease.

Finally, our observations are aligned with predictions from current models of sex-chromosome evolution (14,38–40). Unlike autosomes, the chrX spends approximately two thirds of its evolutionary time in females, exposing X-linked genes to distinct sex-specific selective pressures. For instance, this could lead to X-linked genes evolving faster in response to female-specific selection pressures. In XY individuals, recessive mutations on the X are immediately exposed to selection since compensation by a second allele is not possible, thus leading to stronger purifying selection against deleterious mutations in males. But if recessive X-linked alleles confer a male-specific advantage, this would be of immediate benefit in males (since they have only one chrX) and lead to rapid spread in the population, even if such alleles are rare and detrimental to females (since they have two chrX). Once frequent in the population, there would be selective pressure to limit the expression of such alleles to males; the prediction is therefore an enrichment of male-biased genes on the X chromosome (14,38–40). Our observations of specific enrichment of chrX genes among testis-expressed genes and male-specific SDR-related disorders are consistent with this prediction, as is the high frequency of genes on the X expressed only in male germ cells (2). Moreover, based on the analysis of single-cell transcriptomes and genes associated with male-specific SDR-related monogenic disorders, we found a link between chrX genes and both Sertoli cells and spermatogonia. Recent studies have shown that infertility in individuals with Klinefelter syndrome (47,XXY) is linked to the extra dose of some X-linked genes, to undifferentiated fetal germ cells, and to impaired interactions between Sertoli and germ cells (41–43). Whether the set of genes we found involved in male-specific SDR disorders could have dosage-sensitive functions and be involved in the fertility phenotypes observed in Klinefelter patients will be interesting to explore.

One of the limitations of our study and others before is that we have been considering protein-coding genes only. In developmental disorders, for instance, similar rates of X-linked coding mutations have been found in males and females, despite males being affected 1.4-fold more – indicating that the coding mutations cannot account for the bias (44). Future studies will benefit from considering noncoding-RNA loci as well, the known number of which in the human genome is still growing (45). Moreover, this study is a “time capsule”; our conclusions depend on the currently available datasets. Advent of long-read RNA sequencing could help to detect other levels of involvement for X-linked genes in expression patterns, specific function or diseases. Spatial transcriptomics will also be very valuable to build a more refined view of how “sexy” the human X chromosome is at tissue-scale.

## MATERIALS AND METHODS

### Databases used

The Genotype-Tissue Expression (GTEx) portal (https://www.gtexportal.org/home/) is an open-access resource for exploring tissue- and cell-specific gene expression across non-diseased human tissues and organs. It includes bulk RNA-seq data from approximately 1,000 individuals. The Human Protein Atlas (https://proteinatlas.org) also provides comprehensive gene expression profiles, integrating transcriptomic data (including from GTEx, and single-cell data) with proteomic data based on antibody staining covering normal tissues, pooled for both sexes outside of non reproductive organs. It serves as a valuable tool for analyzing protein-coding gene set enrichment across the human body. To extract the gene count per chromosome for normalisations, we used the website from the HGCN – HUGO Gene Nomenclature Committee (https://www.genenames.org/about/). No data requiring ethical approval for animal welfare nor patient consent were generated, beyond the database mentioned.

### Analysis of gene expression across human tissues

To calculate X:autosome expression ratios (Fig. 1 and S1), we used the file “rna_tissue_consensus.tsv” (“consensus” transcript expression levels) for tissue bulk analysis, downloaded from the Human Protein Atlas webpage (version 23.0) in October 2024; and the file "rna_single_cell_type.tsv" (based on single-cell data) for cell-type analysis, downloaded from the Human Protein Atlas webpage (version 25.0) in December 2025. As defined in the HPA website, the “consensus normalized expression value is calculated as the maximum nTPM value for each gene in the two data sources [HPA and GTEx]; for tissues with multiple sub-tissues (brain regions, lymphoid tissues and intestine) the maximum of all sub-tissues is used for the tissue type”. The “rna_tissue_consensus.tsv” file was used as input for the script “Code_X-A ratios_high-exp_v3.R”; the "rna_single_cell_type.tsv" was used as input for the script “Fig1_sc-analysis_tissue-class.R”. Briefly, only genes for which nTPM or nCPM was equal or above 3 were considered, then each gene was associated with a chromosome and average X:autosome expression ratios were calculated and plotted. To associate genes with chromosomes, we used the BioMart data mining tool: https://www.ensembl.org/info/data/biomart/index.html (46). As a filter, we used the file genes.csv (generated from the script “List-genes”, with the unique gene names from the “rna_tissue_consensus.tsv” file) and exported gene names and chr names; before importing results to R (in the scripts mentioned), we filtered out non-canonical chromosome names. Specific scripts were created to generate each of the panels in Figure 1 (“Plot_Brain.R”, “Plot_Breast.R”, etc).

Lists of ‘elevated expression’ per tissue were downloaded from the Human Protein Atlas webpage (version 23.0) in January/February 2024. As per information available in the website, this dataset includes tissue-enriched genes (at least four-fold higher mRNA level in a particular tissue compared to any other tissues), group-enriched genes (at least four-fold higher average mRNA level in a group of 2-5 tissues compared to any other tissues) and tissue-enhanced genes (at least four-fold higher mRNA level in a particular tissue compared to the average level in all other tissues/cell types); this represents a total of 10930 genes out of 20162 (54%). The full list of genes with elevated expression was downloaded for 36 tissues and gene numbers corresponding to the X chromosome, as well as three autosomes (5, 7 and 16) chosen for their similar gene count to the X, were compiled for making our data set. For single-cell expression, lists of ‘elevated expression’ per cell type were downloaded from the Human Protein Atlas webpage (version 25.0) in December 2025 (26).

To assess whether genes with elevated expression were enriched on specific chromosomes, we compared the observed number of elevated genes per tissue on chromosomes X, 5, 7, and 16 to expected counts derived from the overall distribution of elevated genes across tissues (scripts “HPA_processing.R” for tissue date, and “HPA_processing_single-cell.R” for cell-type data). For each tissue–chromosome pair, a 2×2 contingency table was constructed and tested for deviation from expectation using either Fisher’s exact test (if any expected gene count was <5) or a Chi-squared test (otherwise). Odds ratios were calculated for Fisher’s tests. Resulting *p*-values were corrected for multiple testing using the false discovery rate (FDR) method.

### Analysis of annotated gene functions related to sexual differentiation and reproduction

Functional enrichment analysis of X-linked genes (excluding CTA genes, or only CTA genes; list in Supplementary Data, sheet 6) was performed using g:Profiler (https://biit.cs.ut.ee/gprofiler/) with *Homo sapiens* selected as the reference organism. Statistical significance was assessed using the default g:SCS multiple testing correction implemented in g:Profiler, and terms with an adjusted p-value < 0.05 were considered significant. Results were exported from g:Profiler and subsequently processed in R (version 4.5.2) for plotting.

Human genes with annotated functions related to SDR were retrieved from AmiGO 2 (version 2.5.17) by defining ‘Organism’ as ‘Homo sapiens’ and ‘Involved in’ as one of the terms in a list containing SDR-related functions and “control” functions (Supplementary Data). For each term, results were downloaded and analysed via our script “AMIGO-plot.R”. Briefly, gene names were matched to chromosome location using Biomart (47) (Biomart parameters – Database: Ensembl Genes 112; Dataset: Human genes (GRCh38.p14); Filters: Input external references ID list: Gene Name(s), [ID-list specified]; Attributes: Chromosome/scaffold name, Gene name). Only protein-coding genes were considered. The number of genes per chromosome was counted and the proportion to the total number of protein-coding genes per chromosome calculated. The number of protein-coding genes per chromosome was retrieved from the HGNC database (https://www.genenames.org/).

### Analysis of genes associated with monogenic disorders related to sexual differentiation and reproduction

Gene associations with monogenic disorders were retrieved from the Online Mendelian Inheritance in Man database by searching within the clinical synopsis data (May 2024) using an extensive list of terms (Supplementary Data,) related to sexual differentiation and reproduction (SDR). OMIM data was converted into a table (Supplementary Data) using the script “Script-convertOMIM_SDR.R”, adapted from Leitão et al, 2022 (6). For each chromosome, we then calculated the fraction of protein-coding genes that are associated with specific SDR terms. The banded chrX (Fig. 5) is derived from (6) and our data were manually added based on each gene position per cytologically identified bands.

### GO analysis

Gene Ontology and pathway process enrichment analyses were done using metascape (https://metascape.org/gp/index.html#/main/step1). The list of 58 genes associated with male specific SDR were uploaded to the platform; *Homo sapiens* was selected as the reference organism and enrichment analysis was done using the Expression Analysis setting. Metascape identifies significantly enriched pathways and disease associations within the input gene list, integrating Gene ontology biological processes, molecular functions and disease ontology terms. The statistical significance of enrichment is determined by hypergeometric test or Fisher’s exact test, corrected for multiple comparisons, and reported as log10-transformed *p* values. For each enriched term, the number of genes contributing to the enrichment (Count) and the percentage of the total gene list represented (%) are presented. Enriched GO terms and pathway enrichment were ranked based on statistical significance and are summarized in figure 5C.

### Statistical analysis

Each figure legend and/or respective methods section contains details on which statistical tests were used.

## DATA AVAILABILITY

Data, data files and scripts are available through online supplementary material and/or via Zenodo at: https://zenodo.org/records/21612769?preview=1&token=eyJhbGciOiJIUzUxMiJ9.eyJpZCI6ImZiZjVjZWRkLTViZDItNGI2Mi1hODE5LTg3ZDkzYzUxZmNhYiIsImRhdGEiOnt9LCJyYW5kb20iOiJmMmM1ZGZmN2I2YTQ5ZGEzYzI2NmU5YzNiNzdiNjQwNiJ9.JZN04cN7_-zIa1HCBExkW-jW6uGQ32TzkRIPPvmZ39M8bbmq6LE17k43sbT793zs-MmQjnJfRVo6GmcAFGVfkw Accessible to reviewers during peer review. A public DOI will be released upon acceptance.

## AUTHOR CONTRIBUTIONS

K. Ancelin and R. Galupa equally conceived and designed the study, acquired, analysed and interpreted the data, and drafted, revised and approved the current manuscript. P. Somasundaram acquired, analysed and interpreted data, and revised and approved the current manuscript. All authors agree to be accountable for all aspects of the work.

## FIGURE LEGENDS

**Figure S1.**
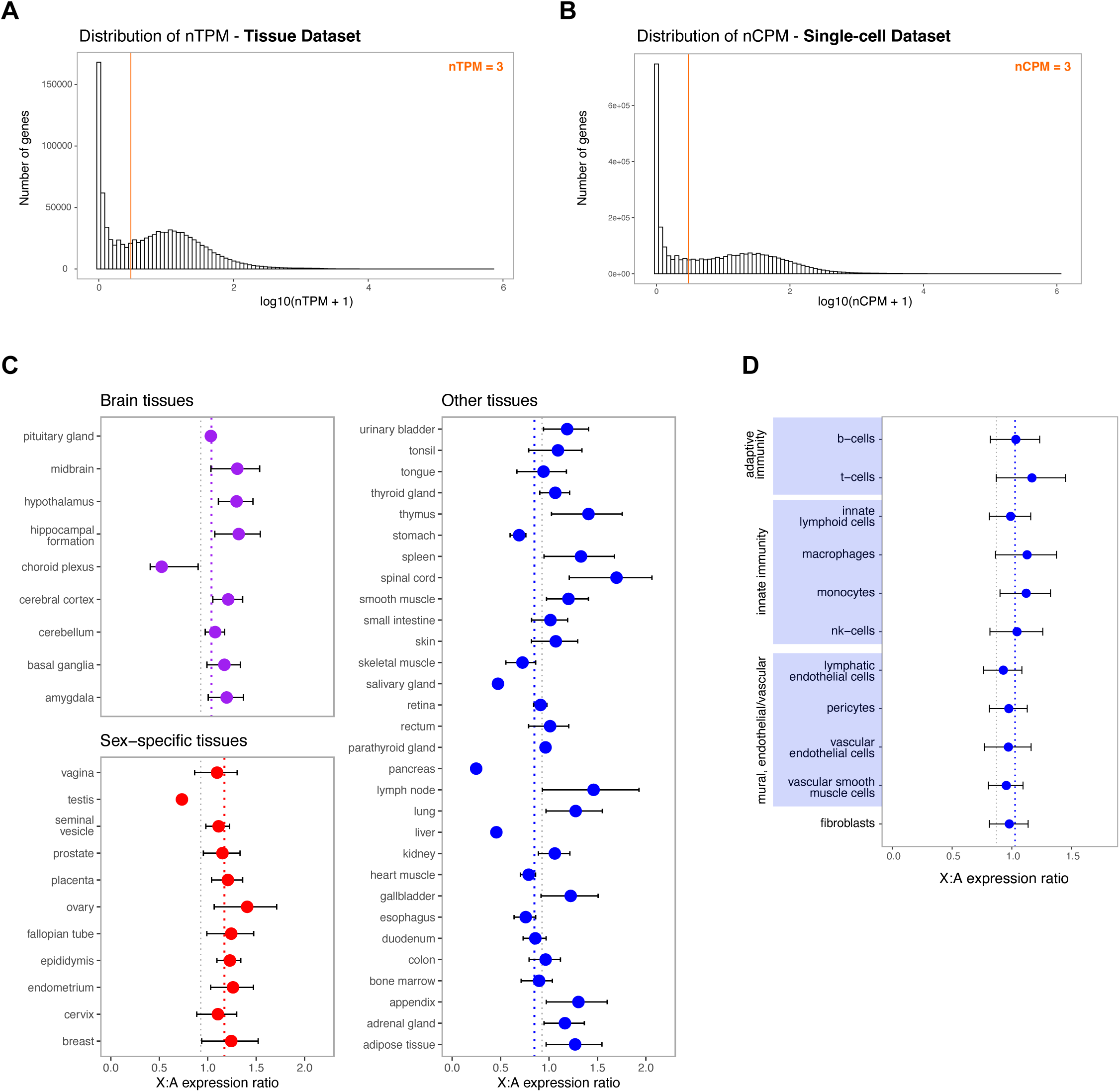
(A-B) Histograms showing the distribution of transcript abundance values for ∼20.150 protein-coding genes plotted as log10(nTPM or nCPM + pseudocount of 1) using 100 bins, based on the “consensus” transcriptomic data from 50 human tissues (A) and 154 cell types (B). The coloured vertical green line represents the nTPM or nCPM threshold to define‘expressed’ genes (nTPM or nCPM=3) for subsequent analyses. (C) X:autosome expression ratios for human adult brain tissues (top left), other somatic tissues (right), and sex-specific tissues (bottom left). Each point represents the mean X:autosome expression ratio (± s.e.m.) for a given tissue listed at the left of each panel. Coloured dotted lines represent the average X:autosome expression ratio for each of the groups of tissues analysed; grey dotted line represents the overall average X:autosome expression ratio (∼0.93). (D) X:autosome expression ratios for cell types related to adaptive immunity (top), innate immunity (second), mural or endothelial cells and fibroblast cells (undefined tissues) (bottom). Each point represents the mean X:autosome expression ratio (± s.e.m.) for a given cell type listed on the left. The coloured dotted line represents the average X:autosome expression ratio for the group of cell types analysed. The grey dotted line represents the overall average X:autosome expression ratio for the single-cells dataset (∼0.87).

**Figure S2.**
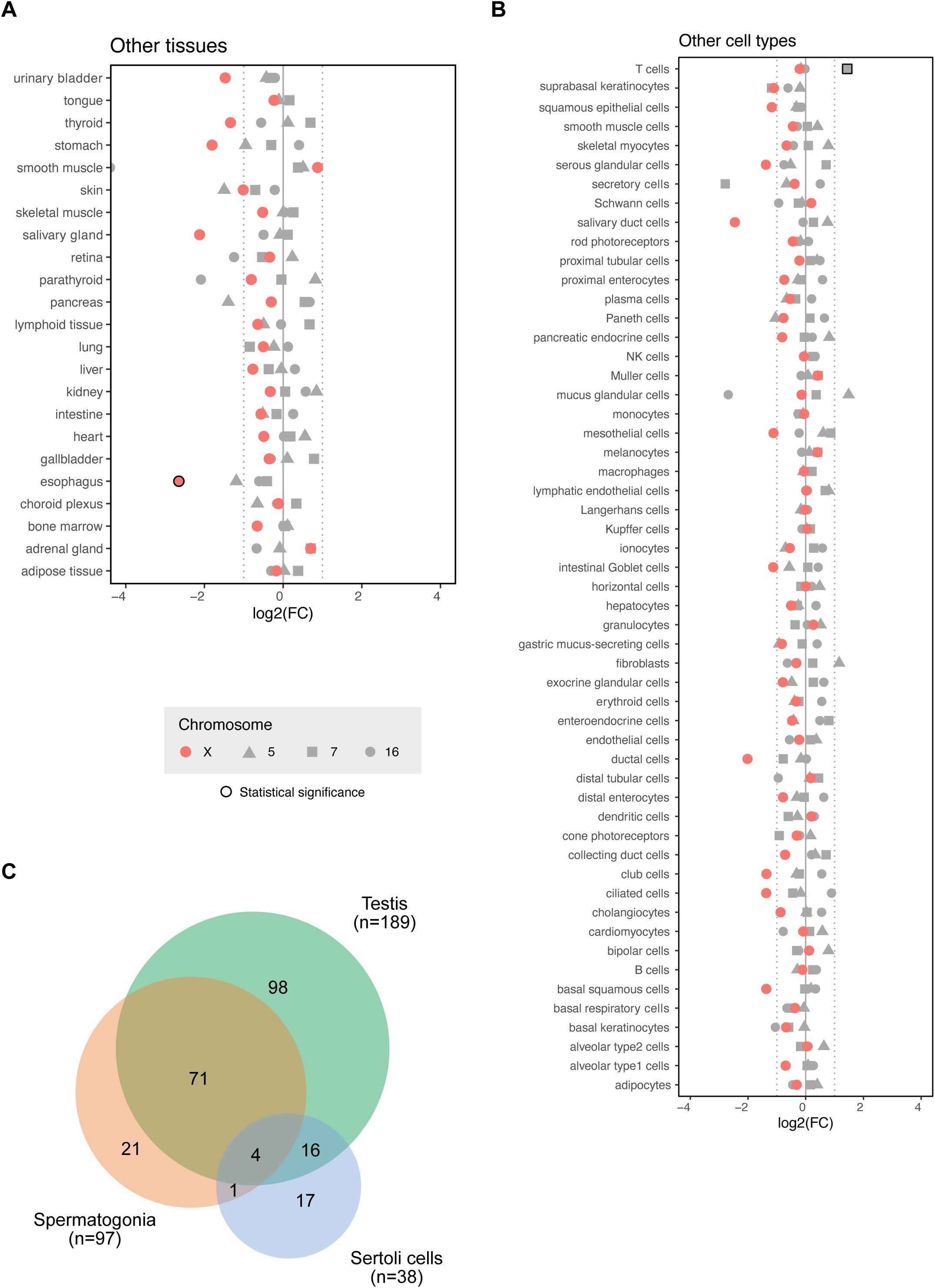
(A-B) Dot plots for log2-transformed ratio (fold-change, FC) between observed and expected frequency for genes of four tested chromosomes among the “elevated” group of genes for “other tissues” (A) and “other cell types” (B). Each chromosome (X, 5, 7 and 16) is represented by a different symbol (respectively an orange circle, a triangle, a square and a grey circle). Statistical test: Chi-squared or Fisher’s test (see Materials and Methods); significance (*p*-value < 0.05) is represented as a black outline around the respective symbol (see Supplementary Data, sheet 2 for *p*-values). Multiple testing correction was performed using false discovery rate (FDR). (C) Venn diagram comparing chrX genes with “elevated expression” in testis, in spermatogonia and in Sertoli cells. Number of respective overlapping genes are indicated.

**Figure S3.**
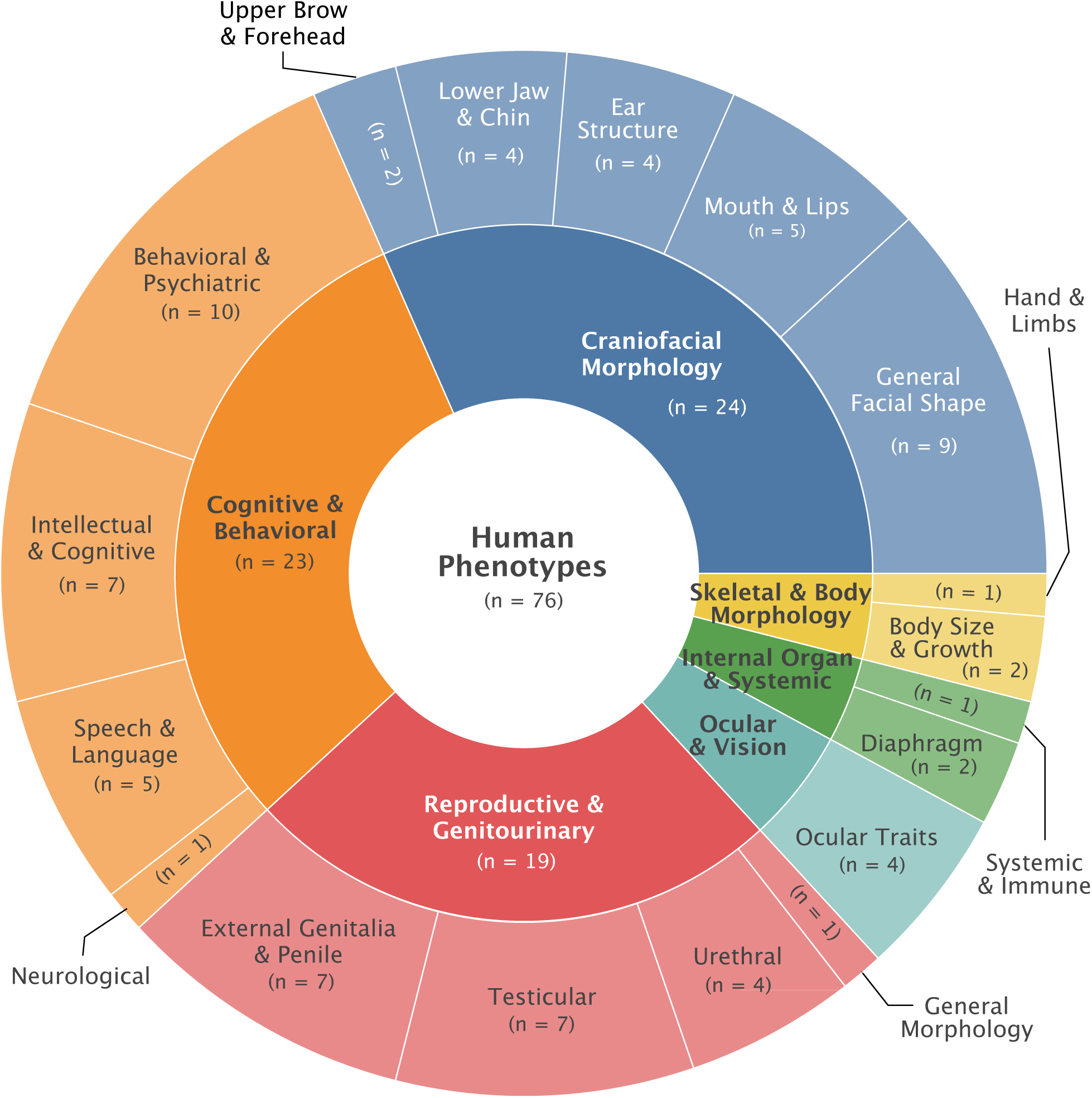
Sunburst plot showing the hierarchical classification of enriched Human Phenotype (HP) terms identified by g:Profiler. Individual HP terms (Supplementary data sheet 6) were manually grouped into higher-order phenotypic categories according to the primary affected organ system or biological function.

## ACKNOWLEDGEMENTS

The authors would like to thank: Catherine Patrat and Noémie Ranisavljević for helpful discussions regarding the medical terms used for the OMIM searches; Elsa Leitão and Christel Depienne for help with their published scripts; and Julie Chaumeil and members of the Galupa team for critical reading of the manuscript. The authors acknowledge the use of ChatGPT for help with coding.

