## Supplementary material for "Reassessing the “sexy” X hypothesis reveals distinct cellular and disease signatures of X-linked genes in male sexual differentiation and reproduction": Table 1

Table 1 – Conclusions of studies addressing gene content on the human and/or mouse X chromosome related to sexual differentiation and reproduction

| Authors, Year | Species | Summary of findings |
|---|---|---|
| Saifi and Chandra, 1999 (20) | Human | The authors analysed how many genes were reported with mutation(s) affecting SDR using both OMIM and GENDIAG databases and manually curated annotations. The authors observed that, compared to autosomes, the X is significantly enriched in SDR-related genes. Results are for both sexes; sex was not analyzed as a variable. |
| Wang et al. 2001 (21) | Mouse | The authors isolated 25 genes that were expressed in spermatogonia but not in somatic tissues and noticed that 10 of them were X-linked, representing a number at least five times bigger than expected by random distribution. |
| Lercher et al. 2003 (27) | Human | The authors determined the percentage of X-linked genes among genes only expressed in prostate, or only in ovaries, or only in mammary gland (based on expression data from SAGE). The authors observed an enrichment for X-linked genes among prostate-only genes, but not among ovary-only or mammary-only genes. |
| Khil et al. 2004 (28) | Mouse | The authors analysed previously published microarrays (Affymetrix) for 49 tissues and found (i) an enrichment for X-linked genes among the most highly expressed genes in ovary and in placenta, and (ii) a depletion of X-linked genes among most highly expressed genes in testis, specifically in post-meiotic stages. |
| Ross et al. 2005 (23) | Human | Upon assembling the human X chromosome sequence, the authors observed that 32 genes on the X contain the MAGE protein domain, which characterises CTA genes, compared to only 4 on the rest of the genome. |
| Hofmann et al. 2008 (34) | Human | By combining expression information from several published datasets, the authors analysed CTA gene expression profiles across several tissues, including ovaries and testis. They observed that CTA genes are predominantly located on the X chromosome and that X-linked CTA genes are significantly more testis-restricted than CTA genes located on autosomes. |
| Mueller et al. 2008 (25) | Mouse | The authors analysed the expression of 33 multicopy gene families on the mouse X (representing ~273 genes),across several tissues, including ovaries and testis, based on microarray data and RNA FISH. They observed that expression of X-linked genes in testis, including at postmeiotic stages, was comparable to autosomal genes. |
| Zhang et al. 2010 (24) | Human and Mouse | The authors analysed expression data from microarrays of several human and mouse tissues. They observed that evolutionary younger X-linked genes were enriched among genes expressed in male post-meiotic stages. In contrast, evolutionary older X-linked genes were depleted. These older X-linked genes showed higher expression in ovaries than autosomal genes. |
| Deng et al. 2011 (26) | Human and Mouse | Based on several RNA-sequencing datasets of human and mouse tissues, including testis and ovaries, the authors found that testis (and brain) have the highest percentage of expressed X-linked genes (84% testis, 78% brain, 23-40% other tissues) |
| Julien et al. 2012 (29) | Human | Based on RNA-sequencing datasets of several human tissues (ovaries not included), the authors observed that the proportion of testis-specific genes is larger for the X than for autosomes. |
